# T cell transcriptomes in simulated microgravity *in vivo*

**DOI:** 10.64898/2026.11.01.763488

**Authors:** Nik V. Kuznetsov

## Abstract

Spaceflight exposes astronauts to a combination of stressors, among which microgravity remains the least well characterized at the level of human immune cell function. We examined the *in vivo* effect of long-term simulated microgravity (SMG) on T cell transcriptomes in ten healthy male volunteers who completed a three-week dry immersion protocol without countermeasures. Peripheral blood was collected at five time points covering a pre-immersion baseline (day -7), three intra-exposure time points (days 7, 14 and 21) and a recovery time point one week after the end of exposure (day 28). CD3+ T cells were isolated and analysed by RNA-seq, with differential expression assessed per volunteer and KEGG pathway enrichment used to identify the most affected gene networks. SMG was associated with transcript-level changes in genes mapping to key molecular pathways, including those involved in cell proliferation, cell-cycle regulation, maintenance of genome integrity, and oncogenic and tumour-suppressor networks. Hub genes shared across the most significant pathways were differentially expressed in the majority of volunteers. A notable finding was pronounced inter-individual heterogeneity: samples clustered primarily by volunteer identity, yet a coordinated temporal shift during exposure and a partial return toward baseline by day 28 were detectable across donors, and these changes occurred in the absence of detectable systemic inflammation. These results indicate that SMG is associated with altered T cell transcriptional programs *in vivo* and that the response is sufficiently variable between individuals to favor personalized rather than uniform countermeasure strategies.

## 1. Introduction

The spaceflight environment presents a complex spectrum of physiological stressors that profoundly differ from conditions on Earth, posing significant risks to astronaut health and mission success. NASA has categorized these into five primary hazards: isolation, confinement, distance from Earth, gravity alterations, and exposure to space radiation [1, 2]. Furthermore, additional factors such as hypomagnetic environment, circadian rhythm disruption, and perturbations in the environmental microbiome can further compound these challenges and influence crew wellbeing [3]. Among these stressors, the effects of galactic cosmic radiation (GCR) on biomolecular and cellular processes have been extensively characterized through both spaceflight and ground-based simulation studies [4, 5]. In contrast, the specific contribution of microgravity to cellular dysfunction, particularly in immune cell signaling, proliferation, and metabolism, remains insufficiently defined and a subject of active investigation [6]. Recent evidences suggest that MG may be a dominant driver of the physiological stress observed in spaceflight, significantly impacting biological systems [7, 8].

To delineate the specific *in vivo* impacts of microgravity on human T lymphocytes, we applied long-term SMG to healthy volunteers. Our analysis of T cell transcriptomes revealed significant deregulation of core gene networks and oncogenic signaling pathways. A critical and novel finding was the observation of substantial inter-individual heterogeneity, with several distinct types of transcriptional responses identified among the volunteers. This evidence compellingly supports a shift in perspective: rather than pursuing a universal solution, effective countermeasures may require a personalized medicine approach that accounts for individual variability in response to microgravity.

A microgravity investigation, conducted on a cohort of ten healthy individuals, was designed to determine whether the absence of gravity alone elicits detectable changes of T cell transcriptomes *in vivo*. Our findings could not only shed light on fundamental mechanisms of space cell biology but also serve as a pivotal stepping stone toward developing targeted strategies to safeguard human health during extended stays in microgravity.

## 2. Methods

### 1.1. Simulated microgravity experiment setup

Ten healthy male participants, aged 24 to 32 years, participated in a three-week simulated microgravity (SMG) experiment using dry immersion, with no countermeasures administered, as outlined in [7]. All volunteers gave their written consent prior to the study, adhering to the ethical requirements of the Helsinki Code of Medical Ethics regarding Human Specimens.

### 1.2. Peripheral blood sample collection

During the SMG protocol, peripheral blood samples (10 mL each) were collected from every volunteer into sodium heparin tubes at five predefined time points: 7 days prior to SMG onset (day –7; background, bg), at 7, 14, 21 days of microgravity exposure, and 7 days after the end of SMG exposure (day 28) (Figure 1).

**Figure 1.**
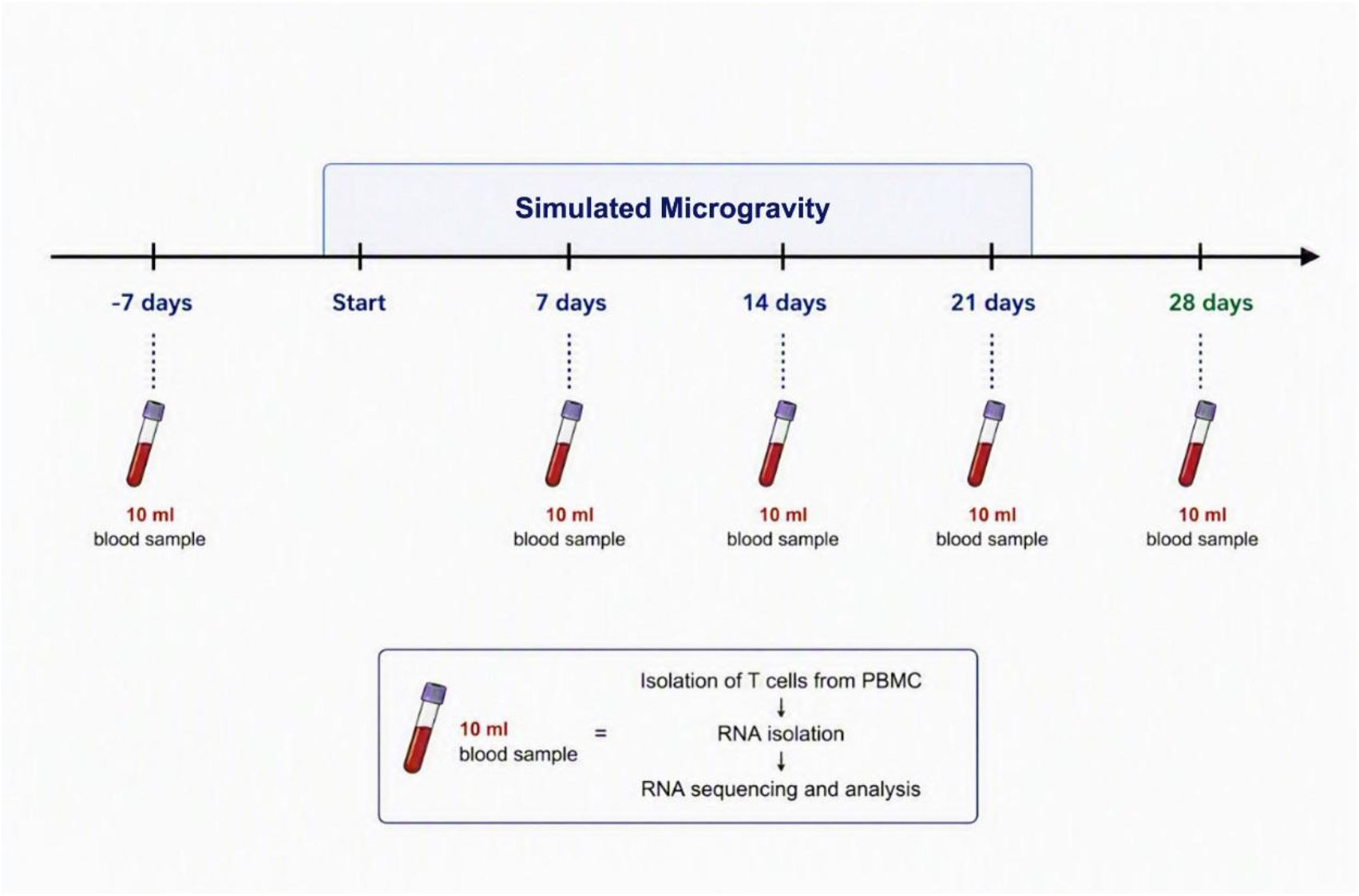
Procedural setup for the ground-based SMG study. The time course of the SMG experiment and sample processing flow chart.

### 1.3. CD3+ T cell isolation

Ficoll-Paque™ PLUS (Cytiva) was employed to obtain PBMCs. CD3+ T cell subset was then positively selected (Stem Cell Technology), resulting in a typical purity of 95%.

### 1.4. RNA Extraction and Sequencing

RNA was extracted, quantified, and RNA quality was assessed via AGE in 1% UltraPure Agarose (Cat. #16500500; Invitrogen, Waltham, MA, USA) as described previously [7]. Gel imaging conducted on a ImageQuant LAS 4000 imaging system as described [9]. Following successful completion of both internal and external quality control assessments (Novogene, Hong Kong), the RNA libraries were processed to 150bp paired-end PE sequencing (Illumina HiSeq; Novogene), followed by standard data quality control procedures.

### 1.5. RNA-Seq Bioinformatics

Sequencing reads mapped to GRCh38/hg38 genome assembly employing HISAT2 (v.2.0.5). Initial bioinformatic preprocessing (quality filtering, alignment, and GO term enrichment) was conducted by Novogene (Hong Kong). Differential gene expression was assessed using the DESeq2 framework within R. To identify expression patterns, log_2_-transformed ratios were clustered via hierarchical clustering, K-means partitioning, and self-organizing maps (SOM). PCA and heatmaps were generated in NASQAR (NYUAD). Functional annotation of genes associated with the identified peaks *was carried out employing* the NCBI Gene portal as detailed previously [6, 10, 11].

## 3. Results

### 1.1. RNA-seq and gene expression data

A total of fifty samples (10 volunteers × 5 time points) yielded up to 58,676 transcripts expression data, which were assigned to over 200 cell signaling pathways. RNA-seq dataset is publicly accessible via the NCBI GEO database under accession number GSE301964.

Differential expression analysis was performed for T cell samples at each time point, with background expression values derived from the pre-immersion baseline (−7d). Transcript levels of differentially expressed genes (DEGs) were visualized on volcano plots of −log10 (q-value) vs. log2(FC) for each time point.

#### 1.2. Gene networks and pathways

To identify the most prominently affected cell signaling networks, twelve KEGG pathways with the highest statistical significance (lowest corrected p-values) were selected from the complete pathway annotation list (Table 1).

**Table 1.**
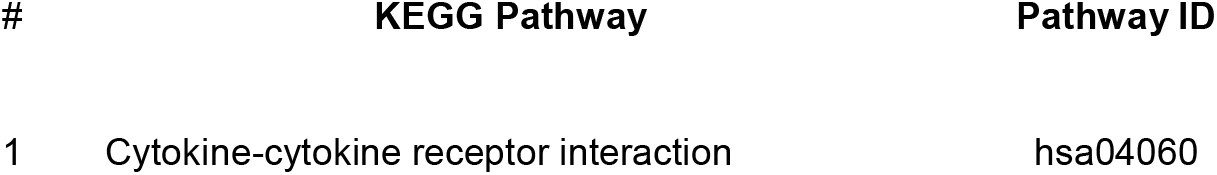
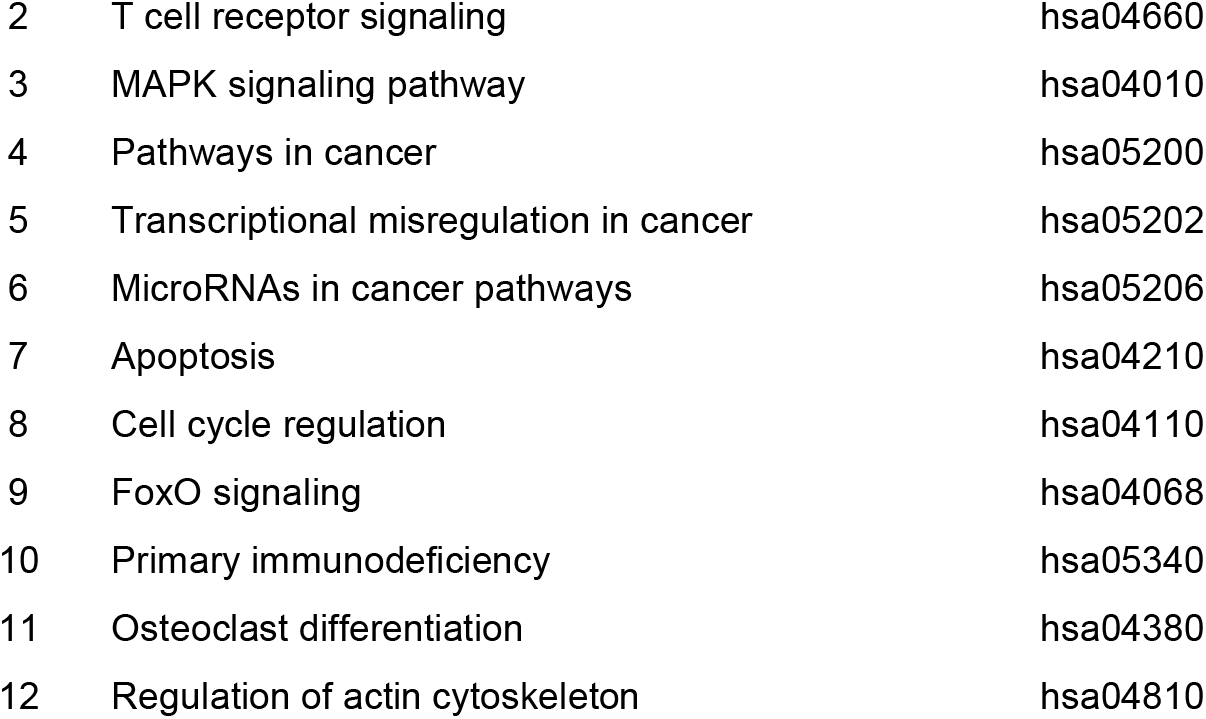
Top 12 KEGG pathways with the highest statistical significance.

| # | **KEGG Pathway** | **Pathway ID** |
|---|---|---|
| 1 | Cytokine-cytokine receptor interaction | hsa04060 |
| 2 | T cell receptor signaling | hsa04660 |
| 3 | MAPK signaling pathway | hsa04010 |
| 4 | Pathways in cancer | hsa05200 |
| 5 | Transcriptional misregulation in cancer | hsa05202 |
| 6 | MicroRNAs in cancer pathways | hsa05206 |
| 7 | Apoptosis | hsa04210 |
| 8 | Cell cycle regulation | hsa04110 |
| 9 | FoxO signaling | hsa04068 |
| 10 | Primary immunodeficiency | hsa05340 |
| 11 | Osteoclast differentiation | hsa04380 |
| 12 | Regulation of actin cytoskeleton | hsa04810 |

From the DEGs mapping to these twelve top-ranked pathways, 44 interconnected genes that appeared in at least two of the selected pathways were designated as hub genes. These genes were subsequently subjected to detailed comparative expression analysis across all time points (Table 2).

**Table 2.** 44 differentially expressed hub genes found in top 12 KEGG pathways.

| Ensemble ID | Gene | Gene Name | Gene ID |
|---|---|---|---|
| ENSG00000125538 | IL1B | interleukin 1 beta | 3553 |
| ENSG00000168685 | IL7R | interleukin 7 receptor | 3575 |
| ENSG00000121966 | CXCR4 | C-X-C motif chemokine receptor 4 | 7852 |
| ENSG00000169429 | CXCL8 | C-X-C motif chemokine ligand 8 | 3576 |
| ENSG00000182578 | CSF1R | colony stimulating factor 1 receptor | 1436 |
| ENSG00000100368 | CSF2RB | colony stimulating factor 2 receptor subunit beta | 1439 |
| ENSG00000119535 | CSF3R | colony stimulating factor 3 receptor | 1441 |
| ENSG00000232810 | TNF | tumor necrosis factor | 7124 |
| ENSG00000067182 | TNFRSF1A | TNF receptor superfamily member 1A | 7132 |
| ENSG00000121858 | TNFSF10 | TNF superfamily member 10 | 8743 |
| ENSG00000163600 | ICOS | inducible T cell costimulator | 29851 |
| ENSG00000177606 | JUN | Jun proto-oncogene, AP-1 transcription factor subunit | 3725 |
| ENSG00000170345 | FOS | Fos proto-oncogene, AP-1 transcription factor subunit | 2353 |
| ENSG00000107968 | MAP3K8 | mitogen-activated protein kinase kinasekinase 8 | 1326 |
| ENSG00000100906 | NFKBIA | NFKB inhibitor alpha | 4792 |
| ENSG00000077150 | NFKB2 | nuclear factor kappa B subunit 2 | 4791 |
| ENSG00000145675 | PIK3R1 | phosphoinositide-3-kinase regulatory subunit 1 | 529 |
| ENSG00000010671 | BTK | Bruton tyrosine kinase | 695 |
| ENSG00000130522 | JUND | JunD proto-oncogene, AP-1 transcription factor subunit | 3727 |
| ENSG00000136997 | MYC | MYC proto-oncogene, bHLH transcription factor | 4609 |
| ENSG00000116717 | GADD45A | growth arrest and DNA damage inducible alpha | 1647 |
| ENSG00000099860 | GADD45B | growth arrest and DNA damage inducible beta | 4616 |
| ENSG00000130222 | GADD45G | growth arrest and DNA damage inducible gamma | 10912 |
| ENSG00000175197 | DDIT3 | DNA damage inducible transcript 3 | 1649 |
| ENSG00000066336 | SPI1 | Spi-1 proto-oncogene | 6688 |
| ENSG00000213281 | NRAS | NRAS proto-oncogene, GTPase | 4893 |
| ENSG00000128340 | RAC2 | Rac family small GTPase 2 | 5880 |
| ENSG00000115415 | STAT1 | signal transducer and activator of transcription 1 | 6772 |
| ENSG00000141510 | TP53 | tumor protein p53 | 7157 |
| ENSG00000124762 | CDKN1A | cyclin dependent kinase inhibitor 1A | 1026 |
| ENSG00000111276 | CDKN1B | cyclin dependent kinase inhibitor 1B | 1027 |
| ENSG00000100985 | MMP9 | matrix metallopeptidase 9 | 4318 |
| ENSG00000115414 | FN1 | fibronectin 1 | 2335 |
| ENSG00000005961 | ITGA2B | integrin subunit alpha 2b | 3674 |
| ENSG00000073756 | PTGS2 | prostaglandin-endoperoxide synthase 2 | 5743 |
| ENSG00000186350 | RXRA | retinoid X receptor alpha | 6256 |
| ENSG00000172115 | CYCS | cytochrome c, somatic | 54205 |
| ENSG00000169896 | ITGAM | integrin subunit alpha M | 3684 |
| ENSG00000148516 | ZEB1 | zinc finger E-box binding homeobox 1 | 6935 |
| ENSG00000170458 | CD14 | CD14 molecule | 929 |
| ENSG00000140379 | BCL2A1 | BCL2 related protein A1 | 597 |
| ENSG00000143384 | MCL1 | MCL1 apoptosis regulator, BCL2 family member | 4170 |
| ENSG00000185338 | SOCS1 | suppressor of cytokine signaling 1 | 8651 |
| ENSG00000185950 | IRS2 | insulin receptor substrate 2 | 866 |

### 1.3. Cancer genes expression patterns and oncogenic pathways

A panel of DEGs was mapped to three cancer-related KEGG pathways that were among the top twelve most statistically significant pathways: hsa05200, hsa05206, and hsa05202.

Among the 34 detected DEGs mapping to the KEGG Pathways in cancer (hsa05200), 24 genes qualified as hub genes based on their presence in at least two of the top twelve pathways. Non-normalized expression values for each of the 24 hub genes were analyzed individually for each of the 10 volunteers across all time points (Figure 2) with the aligned expression patterns.

**Figure 2.**
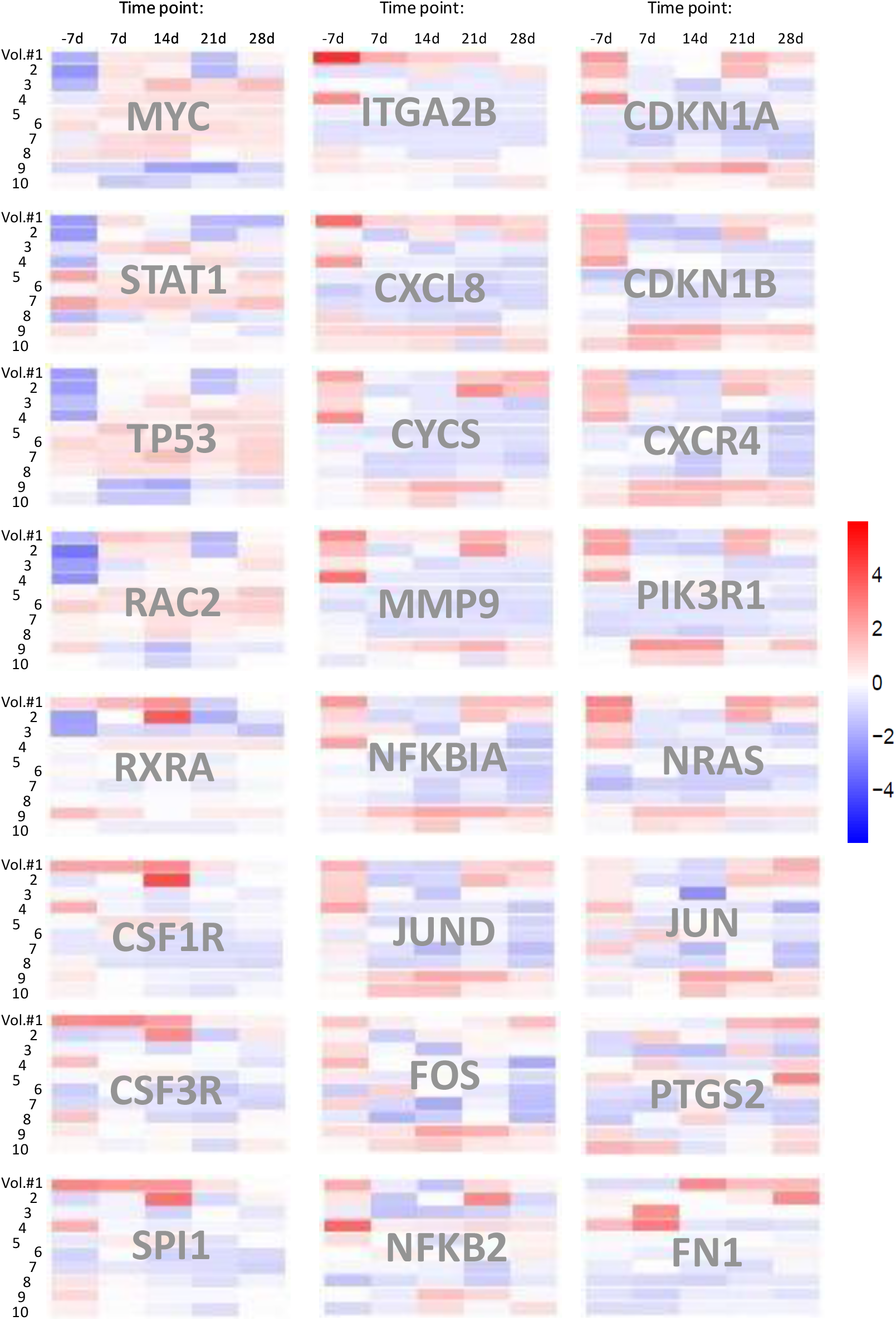
Expression of 24 hub genes in KEGG Pathways in cancer (hsa05200). Non-normalized expression values (Log2 fold change) of detected hub genes were analyzed for each of 10 volunteers (Vol.#1-10) across all time points, and the aligned patterns are shown as heatmaps.

### 1.4. Gene co-expression

Gene co-expression data were analyzed for each volunteer across all time points, and the results are shown as individual Venn diagrams for each of the 10 subjects (Figure 3). Each diagram’s petals quantify time-point-specific genes, with overlaps denoting shared genes across two or more time points.

**Figure 3.**
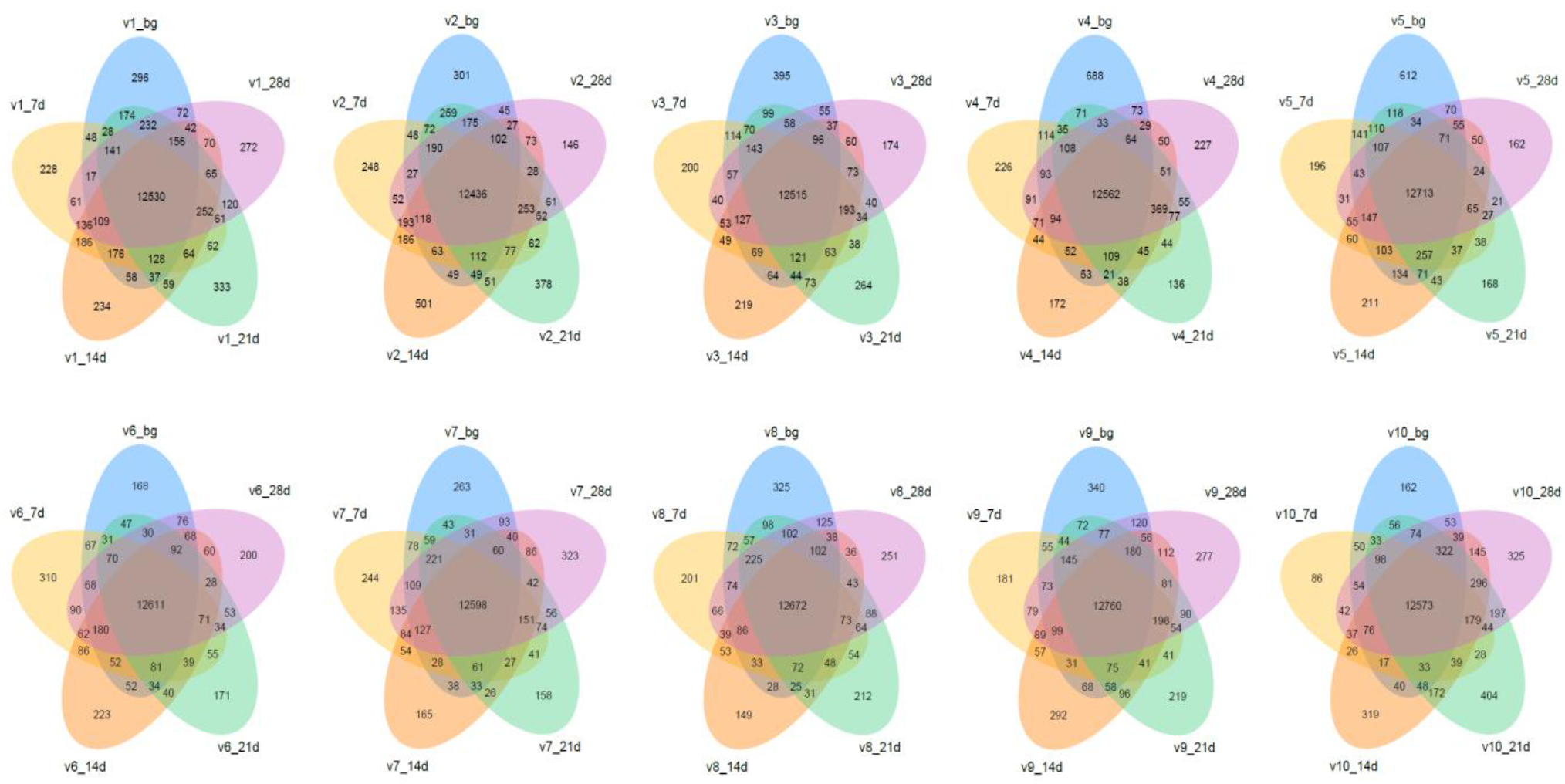
Co-expression Venn diagrams of differentially expressed genes across five time points for each volunteer. Petals: unique gene counts per time point. Overlaps: gene sets shared between ≥2 time points. The sum of the numbers within each ellipse represents the total number of DEGs detected at that time point.

### 1.5. Principal component analysis (PCA)

PCA of the T cell transcriptomic data revealed that samples clustered primarily by volunteer identity rather than by exposure stage across all time points. Volunteers 3–8 consistently formed the main central cluster, whereas Volunteer 9 occupied a distinct and highly separated region of the PCA space throughout the study; Volunteers 1, 2, and 10 were also positioned outside the main cluster but remained comparatively closer to it. Despite this pronounced stratification by individual, coordinated shifts in PCA coordinates were observed between baseline (-7 days) and the immersion period (7–21 days) for all participants. The most pronounced displacements occurred at 14 and 21 days of exposure, while the 28-day samples partially reverted toward their respective pre-imersion positions (Figure 4).

**Figure 4.**
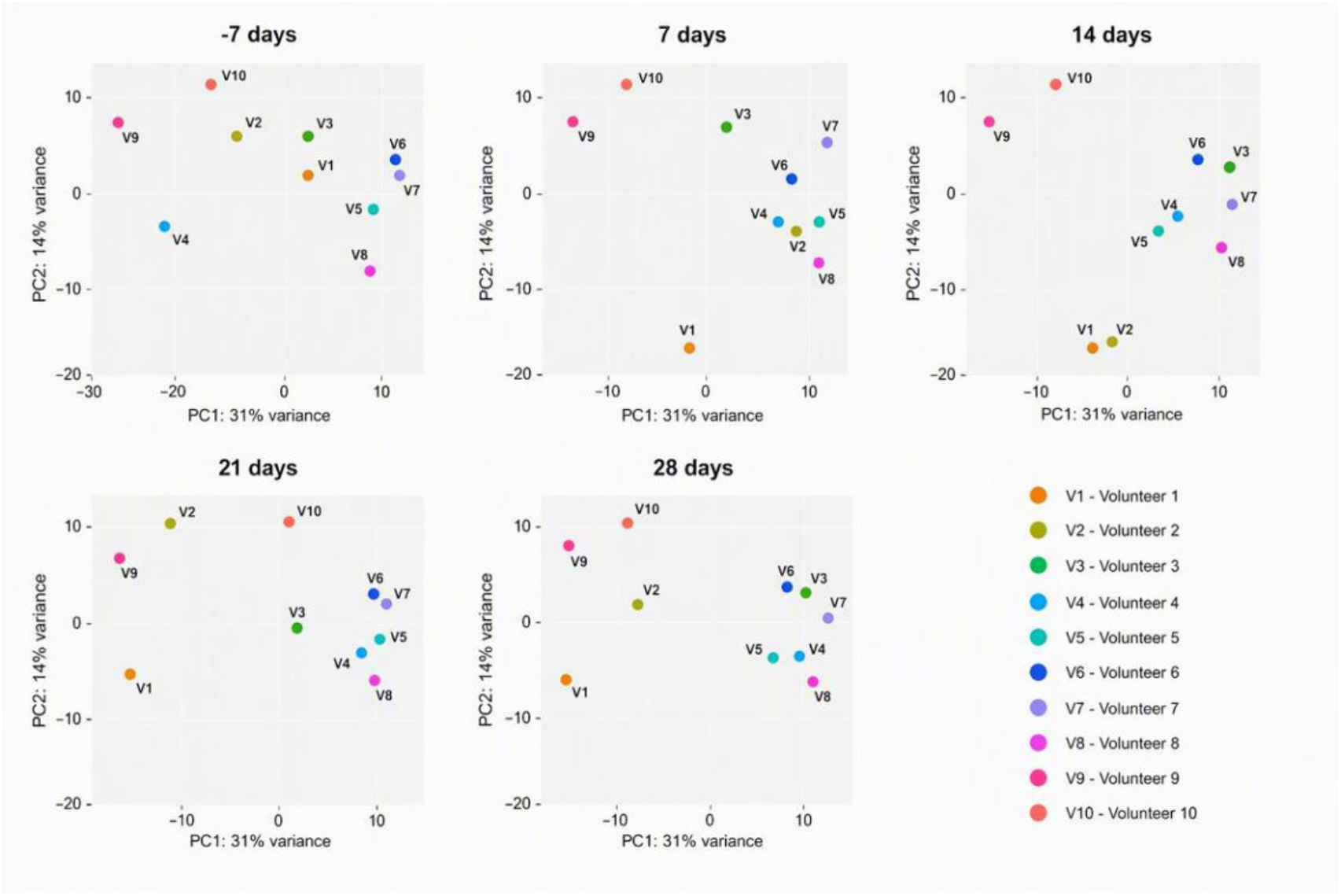
Group principal component analysis (PCA). PCA of transcriptomic profiles across the SMG exposure time course, with each volunteer’s trajectory revealing pronounced inter-individual stratification and heterogeneous response dynamics.

## 4. Discussion

### 4.1. Cytokine-cytokine receptor interaction

The identification of ten genes (IL1B, IL7R, CXCR4, CXCL8, CSF1R, CSF2RB, CSF3R, TNF, TNFRSF1A, TNFSF10) within the KEGG Cytokine-cytokine receptor interaction pathway (hsa04060) reveals a coordinated perturbation of ligand-receptor signaling networks governing inflammation, leukocyte trafficking, and hematopoietic regulation. Microgravity consistently dysregulates cytokine networks and immune cell function, altering expression of pro-inflammatory IL-1β and TNF while shifting inflammatory set points that compromise host defense [12-16]. Recent single-cell analyses have further confirmed that immune suppression signatures are invariant whether evoked by SMG or orbital microgravity, reinforcing the robustness of these findings [17]. Chemokine components CXCL8 and CXCR4 likely mediate microgravity-sensitive changes in neutrophil and lymphocyte chemotaxis and tissue homing, mechanisms repeatedly disrupted in both real and SMG experiments [13, 15]. Colony stimulating factor receptors (CSF1R, CSF2RB, CSF3R) and IL7R directly link to altered myeloid and lymphoid differentiation and hematopoietic output observed in spaceflight studies [13, 16]. TNF, TNFRSF1A, and TNFSF10 highlight plausible routes through which microgravity may modulate apoptosis, survival signaling, and inflammatory amplification via receptor-mediated cascades, providing a molecular framework connecting transcriptional findings to immune and hematopoietic phenotypes documented in experimental literature [12, 14, 18]. The involvement of NF-κB-mediated transcription further underscores the gravity-sensitive nature of these inflammatory pathways [19].

### 4.2. T cell receptor signaling

Eight genes (ICOS, JUN, FOS, MAP3K8, NFKBIA, NFKB2, TNF, PIK3R1) mapping to the T cell receptor signaling pathway (hsa04660) indicate coordinated disruption of antigen receptor-proximal signaling, co-stimulation, transcriptional activation, and downstream survival programs under microgravity. TCR engagement normally recruits MAP kinases and NF-κB while activating AP-1 (JUN/FOS) to drive cytokine production and proliferation; perturbation of these nodes blunts T cell activation and effector differentiation, consistent with gravity-sensitive signaling in lymphocytes [17, 20]. Gravitational mechanosignals governing T cell activation have been molecularly defined, with early studies demonstrating that microgravity inhibits immediate early gene transcription through disruption of the Rel/NF-κB pathway *19, 21+. PI3K regulatory subunits such as PIK3R1 function as central hubs linking mechanotransduction to metabolic and trafficking responses in T cells, offering a mechanistic route through which altered mechanical forces reprogram activation thresholds and migratory behavior [18, 22, 23]. Exposing primary human T cells to both transient and prolonged gravitational unloading, via parabolic flight and clinorotation, respectively, has confirmed the sensitivity of these pathways to altered gravity [24].

Co-stimulatory molecules like ICOS and MAP3K8 further shape TCR signaling amplitude, while changes in NF-κB components (NFKB2, NFKBIA) and TNF can shift the balance between proliferation, survival, and apoptosis under stress [20, 21]. Collectively, these genes suggest that microgravity impairs adaptive immunity by simultaneously dampening co-stimulatory input, rewiring MAPK/PI3K mechanosensitive signaling, and altering NF-κB/AP-1 transcriptional programs essential for robust T cell responses.

### 4.3. MAPK signaling pathway

Ten MAPK pathway genes (JUND, JUN, FOS, MYC, GADD45A, GADD45B, DDIT3, MAP3K8, IL1B, TNF) mapping to hsa04010 point to a gravity-sensitive network coupling stress sensing, transcriptional reprogramming, and cell fate decisions. MAP3K8 and related MAP3Ks occupy upstream nodes determining ERK/JNK/p38 activation patterns; systematic analyses show combinatorial MAPK activation can switch outcomes between proliferation and apoptosis, an effect plausibly translating into altered tissue maintenance under microgravity [25]. Early work demonstrated that coordinated signaling via the MAPK network and SRC-kinase is required for the mechano-induced expression of the immediate-early gene EGR-1 and the pro-inflammatory mediator NF-κB, establishing the mechanosensitivity of this pathway *26+. AP-1 components (JUN/FOS/JUND) and MYC act as immediate early transcriptional effectors of MAPK signaling, controlling cell cycle entry, metabolic reprogramming, and survival, programs modulated by mechanical unloading and environmental stressors [27]. Studies in human bone marrow mesenchymal stem cells have confirmed that SMG inhibits cell proliferation and activates MAPK pathway components [28]. Stress-responsive genes including GADD45A/B and DDIT3 integrate DNA damage and ER stress signals with MAPK outputs, providing mechanistic routes to growth arrest and pro-apoptotic phenotypes reported in cells exposed to SMG [29]. Inflammatory mediators IL1B and TNF within this MAPK module links mechanical perturbation to cytokine-driven amplification loops that reshape MAPK dynamics and downstream transcriptional programs [30]. These data support a model wherein microgravity perturbs MAPK network topology and timing via MAP3K nodes, AP-1/MYC transcriptional control, and stress-responsive effectors, thereby altering proliferation, survival, and inflammatory outputs consistent with experimental observations [29].

### 4.4. Pathways in cancer

The 24 genes mapping to Pathways in cancer (hsa05200) including JUN, FOS, MYC, JUND, SPI1, NRAS, RAC2, NFKBIA, NFKB2, STAT1, TP53, CDKN1A, CDKN1B, PIK3R1, MMP9, FN1, ITGA2B, PTGS2, RXRA, CSF1R, CSF3R, CXCL8, CXCR4, CYCS suggest microgravity engages a cancer-relevant network coordinating cell cycle control, stress responses, survival/apoptosis decisions, cytoskeletal and extracellular matrix remodeling, and signaling through RAS/PI3K and inflammatory axes. Alterations in TP53, CDKN1A/B, MYC, and AP-1 components can reprogram proliferation and senescence programs, while changes in CYCS and TNF/IL1 mediators bias cells toward apoptosis or survival depending on context. Recent work demonstrates microgravity perturbs the p53 regulatory network and aging-associated targets, linking altered gravity to DNA damage and stress signaling converging on tumor suppressor pathways [7, 31]. Studies using colorectal cancer organoids have demonstrated that SMG significantly affects tumor growth patterns and drug responses, providing functional validation of these pathway perturbations [32]. Concurrent modulation of NRAS, RAC2, and PIK3R1 indicates rewiring of proliferative and motility signaling that alters cytoskeletal dynamics and promotes three-dimensional spheroid formation or altered adhesion phenotypes observed in microgravity cancer models, with downstream consequences for drug sensitivity and metabolic state through MYC-dependent programs [33-35]. The potential of using space-based investigations to inform cancer research on Earth has been increasingly recognized, with microgravity revealing fundamental aspects of tumor biology [36]. Matrix remodeling genes (MMP9, FN1, ITGA2B) together with chemokine/chemokine receptor pairs (CXCL8/CXCR4) provide mechanistic basis for shifts in invasion, migration, and immune cell interactions under unloading conditions, potentially unmasking oncogenic behaviors *in vitro* while altering chemotherapy responses [34, 35, 37]. Enrichment of cancer pathway genes in our microgravity signature argues that altered gravitational forces reconfigure core regulatory circuits governing proliferation, stress adaptation, and cell-matrix interactions—changes directly relevant to tissue homeostasis, regenerative capacity, and interpretation of cancer cell phenotypes in space biology studies [7, 31, 38].

#### 4.4.1. Transcriptional misregulation in cancer

Twelve genes mapped to Transcriptional misregulation in cancer (hsa05202): ITGAM, ZEB1, RXRA, CSF1R, CD14, TP53, MMP9, CDKN1A, CDKN1B, MYC, BCL2A1, DDIT3 point to a gravity-sensitive axis wherein altered mechanical input transduces into persistent changes in transcriptional programs governing inflammation, epithelial-mesenchymal plasticity, cell cycle control, and survival. Transcription factors and co-regulators including ZEB1 and MYC can rewire lineage and stress response programs, while modulators of chromatin and nuclear signaling (RXRA) and stress-induced effectors (DDIT3/CHOP, BCL2A1) determine whether cells adopt adaptive, senescent, or apoptotic fates. Recent syntheses of microgravity transcriptomics emphasize such transcriptional reprogramming as a hallmark of unloading underlying many downstream phenotypes [39, 40]. Studies on intestinal stem cells and epithelial regeneration have further highlighted the impact of microgravity on tissue-specific transcriptional programs [41]. Innate immune and myeloid regulators (CD14, CSF1R, ITGAM) together with inflammatory mediator MMP9 provide a direct molecular link between transcriptional misregulation and altered immune-stromal crosstalk in microgravity, consistent with reports that gene regulatory networks controlling inflammation and extracellular matrix remodeling are particularly labile under real and simulated microgravity [40, 42]. SMG has been shown to inhibit osteogenic differentiation through downregulation of ZEB1, demonstrating the functional relevance of this transcription factor in unloading responses [43]. Perturbation of tumor suppressor and cell cycle checkpoints (TP53, CDKN1A, CDKN1B) combined with changes in adhesion and matrix remodeling genes explains frequently observed shifts in proliferation, three-dimensional growth, and stressor resistance/sensitivity in microgravity models [44]. Mechanistic work on ZEB1 further demonstrates how EMT-linked transcription factors can promote replication stress tolerance while creating vulnerabilities in DNA damage responses, a duality plausibly operating in unloaded cells to shape long-term tissue outcomes [39, 42]. These observations support a model wherein microgravity drives durable transcriptional misregulation across immune, stress, and developmental programs via nodes identified in hsa05202, linking altered mechanics to changes in tissue remodeling, immune competence, and cell fate decisions central to organismal adaptation during spaceflight.

#### 4.4.2. MicroRNAs in cancer pathways

The mapping of MYC, NRAS, TP53, DDIT3, MCL1, ZEB1, SOCS1, CDKN1A, CDKN1B, IRS2, MMP9, and PTGS2 to the MicroRNAs in cancer pathway (hsa05206) highlights microRNAs as pivotal post-transcriptional regulators that can amplify or buffer microgravity-induced transcriptional shifts, reshaping cell fate, stress responses, and signaling plasticity. MicroRNAs control many observed genes—miR-34/miR-145 families intersect with MYC and TP53 networks, miR-21 and miR-155 modulate inflammatory and survival nodes including MCL1 and SOCS1, and miR-200 family members regulate EMT drivers such as ZEB1, suggesting gravity-dependent changes in miRNA expression can produce coordinated downstream effects on proliferation, apoptosis, EMT, and metabolic signaling [45]. Integrated peripheral-lymphocytic microRNA/mRNA transcriptome co-profiling of astronauts has revealed consistent dysregulation patterns following spaceflight [46]. Recent omics studies of tumor cells and broader microgravity transcriptomic surveys report consistent dysregulation of miRNA networks under real and simulated microgravity, with consequent alterations in cell cycle regulators (CDKN1A/B), pro-apoptotic effectors (DDIT3), and matrix remodeling enzymes (MMP9), collectively influencing spheroid formation, drug sensitivity, and stress tolerance in unloaded cells [39, 47]. MicroRNA expression profiling in SMG-treated human colon cancer cells has identified specific miRNA-mRNA regulatory networks that mediate adaptive responses [48]. These findings suggest a model wherein microgravity perturbs miRNA biogenesis or stability, modulating abundance and activity of key oncogenic and tumor suppressor targets (MYC, NRAS, TP53, IRS2), with this post-transcriptional layer constituting a major determinant of phenotypic plasticity observed in space biology experiments [49]. Integrating miRNA profiling with gene-level signatures will be essential to resolve causal regulatory chains and identify miRNA-based biomarkers or countermeasures mitigating microgravity-induced dysregulation of proliferation, survival, and immune-related pathways.

### 4.5. Apoptosis pathway

Seventeen genes mapped to the Apoptosis pathway (hsa04210): JUN, FOS, NRAS, NFKBIA, TNF, TNFRSF1A, TNFSF10, CYCS, GADD45A/B/G, DDIT3, MCL1, CSF2RB, PIK3R1, TP53, BCL2A1 define a multilayered program through which microgravity can tip the balance between survival and programmed cell death via mitochondrial, death receptor, ER stress, and pro-survival kinase axes. Mitochondrial apoptosis, centrally mediated by cytochrome c release and BCL-2 family regulation (CYCS, MCL1, BCL2A1), exhibits sensitivity to redox changes and mechanical stress, with repeated implication in microgravity-enhanced apoptosis across cell types [50-53]. Recent studies have identified PIEZO1 as a mediator of endothelial cell apoptosis through HMGA2 under SMG, revealing novel mechanotransductive links to cell death [54]. p53 and stress-responsive effectors (TP53, GADD45 family, DDIT3/CHOP) integrate DNA damage and ER stress signals to promote either repair or mitochondrial death pathways; recent mechanistic reviews highlight how altered gravity amplifies these stress signals and mitochondrial vulnerability [52, 55]. Studies in vascular endothelial cells and lymphoblastoid cells have confirmed that SMG enhances apoptosis through the mitochondrial pathway [56, 57]. Concurrently, death receptor signaling (TNF, TNFRSF1A, TNFSF10) and NF-κB modulators (NFKBIA) provide extracellular and transcriptional inputs that can either reinforce apoptosis or engage compensatory survival programs. PI3K regulatory subunits (PIK3R1) and CSF2RB link cytokine-mediated trophic signaling to AKT/ERK survival cascades, offering routes through which microgravity-induced cytokine dysregulation secondarily modulates apoptotic thresholds. Empirical studies in real and simulated microgravity report increased caspase activation, mitochondrial outer membrane permeabilization, and ER stress markers consistent with this network-level model, indicating that coordinated perturbation of mitochondrial effectors, death receptor ligands, stress sensors, and pro-survival kinases captured by hsa04210 plausibly underlies cell type- and context-dependent apoptosis phenotypes observed during unloading [50-53].

### 4.6. Cell cycle regulation pathway

Seven cell cycle genes (CDKN1A, CDKN1B, GADD45A, GADD45B, GADD45G, MYC, TP53) mapping cleanly to the KEGG Cell cycle pathway (hsa04110) point to coordinated disruption of G1/S checkpoint control, DNA damage sensing, and replication stress responses in unloaded conditions. Elevated expression of cyclin-dependent kinase inhibitor CDKN1A/p21 has been repeatedly observed in bone and other mechanically sensitive tissues during spaceflight and SMG, where increased p21 correlates with cell cycle arrest, reduced proliferation, and impaired regenerative capacity. Genetic or pharmacologic modulation of p21 alters these outcomes in rodent flight models, directly linking CDKN1A dynamics to gravity-dependent tissue maintenance [7, 52, 58, 59]. NASA experiments have specifically investigated the role of CDKN1a/p21 in mesenchymal stem cell proliferation and regeneration under microgravity conditions [60]. Studies in human bone marrow mesenchymal stem cells have confirmed that SMG inhibits proliferation by inducing G1 phase arrest through cell cycle regulator modulation [61]. Stress-inducible GADD45 family members act as sensors coupling DNA damage and environmental stress to cell cycle checkpoints and p38/MAPK signaling; their induction in microgravity can enforce growth arrest or facilitate repair depending on context [58, 62]. TP53 further determines whether cells enter transient arrest, senescence, or apoptotic programs [63, 64, 65, 66, 67, 68, 69, 70, 71, 72]. Microgravity-associated perturbations in p53 signaling and MYC-driven metabolic programs have been reported in recent transcriptomic surveys, suggesting altered gravitational loading reprograms the balance between proliferation and protective arrest [7, 33]. Together, these nodes provide mechanistic explanation for frequently observed reductions in proliferation, altered differentiation trajectories, and increased replication stress in cells exposed to real or simulated microgravity, identifying CDKN1A/p21, GADD45s, TP53, and MYC as candidate molecular levers for countermeasures aimed at preserving tissue regenerative potential during long-duration missions.

### 4.7. FoxO signaling pathway

The presence of IL7R, IRS2, and TNFSF10 in our microgravity signature and their mapping to the FoxO signaling pathway (hsa04068) point to a conserved stress-responsive axis through which altered gravitational loading can reprogram cell survival, metabolism, and immune homeostasis. FoxO transcription factors integrate upstream growth factor and insulin/PI3K-AKT inputs—with IRS2 serving as canonical adaptor linking insulin/IGF signaling to AKT and consequently to FoxO phosphorylation and nuclear exclusion—so changes in IRS2 expression or activity under microgravity could shift FoxO localization and downstream programs controlling oxidative stress defenses, autophagy, and metabolic reprogramming [73]. IL7R signaling intersects with FoxO-regulated lymphoid survival and homeostatic proliferation pathways; altered IL-7R dynamics in unloaded conditions may therefore perturb FoxO-dependent maintenance of T and B cell pools [74, 75]. TNFSF10 (TRAIL) engages extrinsic apoptotic pathways modulated by FoxO targets providing a route through which FoxO activity tunes the balance between adaptive stress responses and programmed cell death in microgravity [76]. The broader roles of FOXOs in acute and chronic neurological disorders further underscore their importance as stress-responsive transcription factors [77]. Collectively, these connections suggest microgravity-induced shifts in growth factor signaling (IRS2), cytokine receptor input (IL7R), and death ligand activity (TNFSF10) converge on FoxO transcriptional control to alter redox homeostasis, cell cycle arrest, autophagy, and apoptosis, mechanisms highlighted in studies linking cytokine and insulin-related signaling to FoxO-mediated homeostasis [73, 75, 78].

### 4.8. Primary immunodeficiency

The detection of ICOS, IL7R, and BTK within the Primary immunodeficiency pathway (hsa05340) suggests microgravity impinges on core nodes required for lymphocyte development, co-stimulation, and B cell receptor signaling, potentially producing phenotypes that phenocopy aspects of primary immunodeficiency. IL7R is essential for lymphoid progenitor survival and B/T lineage differentiation; altered IL7R signaling under microgravity could reduce lymphopoiesis and repertoire diversity, as demonstrated in human B cell progenitors where IL-7R activity drives expansion and V(D)J usage patterns [64]. BTK serves as a central transducer of B cell receptor signals required for maturation and antibody responses; perturbation of BTK expression or activity in altered gravity would likely impair humoral immunity and vaccine responsiveness [79]. ICOS provides critical co-stimulatory input for T follicular helper cell function and germinal center responses, so its dysregulation could further compromise B-T cell collaboration and long-lived antibody production [80]. Spaceflight studies have demonstrated persistent alterations in adaptive immunity, including impaired vaccine responses and altered Tcell differentiation [81, 82]. Monitoring studies of T-cell activation and differentiation in microgravity have confirmed the sensitivity of these developmental pathways to gravitational unloading [83]. Together, these three genes provide a mechanistic link between microgravity-induced transcriptional changes and clinically relevant reductions in adaptive immune competence reported in spaceflight and SMG studies [13, 74, 79].

### 4.9. Osteoclast differentiation

Fourteen genes mapped to Osteoclast differentiation (hsa04380): IL1B, CSF1R, JUND, JUN, FOS, SPI1, SOCS1, NFKBIA, NFKB2, STAT1, TNF, TNFRSF1A, PIK3R1, BTK define a coherent signaling module linking inflammatory cues, receptor tyrosine/kinase inputs, transcriptional effectors, and NF-κB/PI3K survival programs to the generation and activation of bone-resorbing osteoclasts. Colony stimulating factor signaling through CSF1R and pro-inflammatory ligands including TNF and IL1B are established drivers of osteoclast lineage commitment and fusion; in microgravity these cytokine axes are frequently dysregulated, providing molecular explanation for enhanced osteoclastogenesis and accelerated bone resorption observed in unloaded models and astronauts [84,85,86]. Spaceflight studies have directly demonstrated that microgravity affects osteoclastogenesis by increasing the RANKL/OPG ratio [87]. Downstream, MAPK AP-1 components (JUN, FOS, JUND) and NF-κB regulators (NFKBIA, NFKB2) cooperate with lineage transcription factors such as SPI1/PU.1 to induce osteoclast-specific programs, while PIK3R1 and BTK integrate trophic and receptor signals to modulate survival, cytoskeletal reorganization, and resorptive activity [79, 88]. Systems biology analyses have revealed coordinated osteoclast and osteoblast responses to modeled microgravity [89]. Negative feedback modulators like SOCS1 and STAT-dependent signaling tune differentiation magnitude and duration; perturbation of these checks in microgravity can shift balance toward excessive resorption [18, 86]. Enrichment of these genes in hsa04380 supports a model wherein microgravity-induced cytokine imbalance, altered CSF1R/BTK/PI3K signaling, and reprogrammed AP-1/NF-κB transcriptional responses converge to promote osteoclast differentiation and activity, mechanisms aligning with documented microgravity-associated bone loss and osteoclastogenic pathways [52, 84, 90].

### 4.10. Regulation of actin cytoskeleton

The Regulation of actin cytoskeleton (hsa04810) enrichment driven by ITGAM, ITGA2B, CD14, FN1, RAC2, NRAS, and PIK3R1, provides a mechanistic bridge between altered gravitational loading and the rapid, cell type-specific reorganization of adhesion, force transmission, and motility characterizing microgravity responses. Integrin-ECM complexes (ITGA2B/ITGAM; FN1) form the primary mechanical linkage to intracellular actin networks and focal adhesions; disruption of this reciprocity under unloading remodels focal adhesion composition and signaling through RAS/PI3K and Rho family GTPases, producing downstream changes in lamellipodial dynamics, phagocytic cup formation, and cell spreading repeatedly observed in immune and stromal cells in SMG [9, 91, 92, 93, 94]. The cytoskeleton, acting through mechanosensitive channels, is now recognized as a sensor of gravitational fluctuations and a primary mechanotransductive mechanism [95]. RAC2 and NRAS coordinate actin nucleation and membrane protrusion; genetic and functional studies demonstrate Rac1/Rac2 essential for immune cell motility and antigen responses, linking RAC2 signal to impaired chemotaxis and altered immune function reported in unloading conditions [10, 11, 57]. PIK3R1 couples integrin and growth factor receptors to actin remodeling effectors (WASP/WAVE, ARP2/3) and metabolic programs sustaining cytoskeletal turnover, offering a plausible route through which changes in PI3K regulatory subunits translate mechanical cues into persistent transcriptional and phenotypic shifts [22, 92, 93]. Nuclear actin regulators and actin-linked transcriptional co-factors further connect cytoskeletal remodeling to gene expression: nuclear WASP protein modulates T lymphocyte transcriptional programs, illustrating how actin regulators directly influence lineage-specific transcriptional outputs relevant to our immune cell signatures [9]. RhoGTPases have been identified as key players in mechanotransduction during microgravity exposure, linking cytoskeletal dynamics to downstream signaling [96]. Comparative transcriptomic work demonstrates responses to SMG are distinct from hydrodynamic stress yet converge on cytoskeletal and adhesion modules, underscoring actin regulatory network as a reproducible gravity-sensitive hub across experimental systems [10, 11, 97, 98]. Dynamic gene expression response studies in human T cells have further confirmed the centrality of cytoskeletal regulation in gravity sensing [6, 7, 10, 11, 97, 98]. Recent bioinformatic and experimental analyses show microgravity-induced cytoskeletal instability correlates with altered focal adhesion signaling, changes in RAC-dependent motility, and emergence of three-dimensional aggregation or spheroid phenotypes—outcomes mechanistically consistent with the identified gene set, with implications for immune competence, wound healing, and tissue architecture during long-duration missions [93, 95, 97]. The cytoskeleton and its role in T cell migration and activation under microgravity conditions has been extensively characterized, providing functional validation of these pathway perturbations [99]. Together, these data support a model wherein integrin-ECM disruption, RAS/PI3K-Rac signaling, and actin-nuclear coupling form a coherent mechanotransductive axis translating reduced gravitational force into altered cell behavior and durable transcriptional reprogramming [10, 11, 98].

### 4.11. PCA-based inter-individual stratification and heterogeneous SMG response trajectories

The PCA trajectories suggest a strong inter-individual effect that is substantially larger than the group-level effect of SMG. At all time points, samples cluster primarily according to volunteer identity rather than according to exposure stage, indicating that baseline transcriptional differences between individuals remain a dominant source of variance. Specifically, volunteers 3-8 consistently formed the main cluster, whereas volunteer 9 remains highly separated and occupies a distinct region of PCA space throughout the study. Volunteers 1, 2 and 10 are also distinct but somewhat closer to the main cluster. Despite this pronounced stratification, a temporal response to SMG is still apparent. Between baseline (-7 days) and exposure (7-21 days), all individuals display coordinated shifts in PCA space, suggesting a common physiological adaptation. The movement is most pronounced during immersion (particularly at 14 and 21 days), whereas at 28 days the samples partially return toward their initial positions, consistent with recovery following cessation of the intervention. However, the magnitude and direction of these trajectories differ among volunteers, indicating substantial heterogeneity in response. Overall, the PCA analysis supports the conclusion that SMG induces a detectable transcriptomic effect, but that the response is strongly modulated by individual-specific factors, leading to marked stratification of the cohort and variable adaptation dynamics between volunteers.

These findings are consistent with human bed-rest transcriptomic studies, which demonstrate that while a core set of genes responds consistently to unloading, baseline expression profiles are highly individual-specific and remain the primary determinant of variance throughout the intervention [100]. This observation aligns with the broader framework proposed for space medicine, wherein omics-based analyses are essential to capture individual molecular profiles and develop tailored countermeasures [101]. Furthermore, the marked heterogeneity observed in our cohort highlights the growing understanding that genetic variability significantly modulate physiological responses to microgravity, supporting the need for a shift from universal to personalized strategies for astronaut health management during long-duration missions [100, 102].

## 5. Conclusions

### 5.1. T cell transcriptomes demonstrate dynamic pathway-specific responses to microgravity

Gene expression profiles and temporal trajectories confirm that T cell transcriptional profiles undergo a dynamic pathway-specific, time-dependent adaptation to microgravity, rather than a static or monotonic shift. Coordinated displacements from background (-7 days) through the SMG period (7-21 days) indicate a shared physiological response that peaks during the middle-to-late exposure phase. The subsequent partial return toward baseline positions by day 28 further supports a recovery trajectory, underscoring that the transcriptomic impact of microgravity is reversible.

### 5.2. T cell transcriptome modulation by microgravity in vivo affects key immune and oncogenic signaling gene networks

In addition to global expression shifts, the PCA-defined variance points to biologically meaningful perturbations in specific regulatory pathways. The involvement of oncogenic and immunoregulatory cascades is particularly noteworthy, as these pathways are critical for maintaining T cell homeostasis and functional integrity. This suggests that microgravity does not simply induce general transcriptional noise but rather has targeted effects on molecular pathways linking immunity, cell growth control, and stress-responsive signaling, potentially with implications for host defense and long-term cellular health.

### 5.3. T cell transcriptome pathway-specific responses to microgravity in vivo are individually variable

Despite the presence of a common temporal signature, the magnitude and direction of these pathway-specific responses are strongly modulated by baseline inter-individual differences. The stratification of the cohort demonstrates that volunteer identity is the dominant source of transcriptional variance. This marked heterogeneity implies that the same microgravity stimulus elicits divergent molecular outcomes across individuals, likely driven by pre-existing baseline transcriptomic profiles. Therefore, studies of T cell adaptation to spaceflight must account for significant interindividual variability, highlighting the need for personalized biomarkers and tailored countermeasures in future space missions.

## Author Contributions

N.K. contributed to the conceptual idea of the paper, formulated the objective, wrote the manuscript, prepared the figures for data presentation, and performed the editing.

## Funding

The study was supported by ASPIRE, the technology program management pillar of Abu Dhabi’s Advanced Technology Research Council (ATRC), via the ASPIRE Precision Medicine Research Institute Abu Dhabi (ASPIREPMRIAD) award grant number VRI-20-10.

## Data Availability Statement

The data that support the findings of this study are currently available in the NCBI GEO public repository.

## Acknowledgments

The author thanks colleagues at the New York University Abu Dhabi, Karo-linska Institute, and Institute of Biomedical Problem for the assistance and discussions.

## Conflicts of Interest

The author declares no conflict of interest.

## Abbreviations

BG: background
MG: microgravity
PBMC: peripheral blood mononuclear cells
PCA: principal component analysis
QC: quality control
SMG: simulated microgravity

## Notes

### Competing Interest Statement

The authors have declared no competing interest.

## References

1. NASA. Risks of Human Spaceflight [Internet]. 2026 [cited 2026 Jul 25]. Available from: https://www.nasa.gov/hrp/risks/

2. Moreno-Villanueva M, Jimenez-Chavez LE, Krieger S, Ding LH, Zhang Y, Babiak-Vazquez A, et al. Transcriptomics analysis reveals potential mechanisms underlying mitochondrial dysfunction and T cell exhaustion in astronauts’ blood cells in space. Front Immunol. 2025;16. doi:10.3389/fimmu.2025.xxxxx

3. Miranda S, Vermeesen R, Janssen A, Rehnberg E, Etlioglu E, Baatout S, et al. Effects of simulated space conditions on CD4+ T cells: a multi modal analysis. Front Immunol. 2024;15:1443936. doi:10.3389/fimmu.2024.1443936

4. Wadhwa A, Moreno-Villanueva M, Crucian B, Wu H. Synergistic interplay between radiation and microgravity in spaceflight-related immunological health risks. Immun Ageing. 2024;21(1). doi:10.1186/s12979-024-00498-9

5. Siddiqui R, Qaisar R, Elmoselhi A, Khan NA. Digital twin modelling in microgravity: A framework for predictive and personalised space medicine. Life Sci Space Res. 2025. doi:10.1016/j.lssr.2025.01.003

6. Kuznetsov NV. Expression of the p53 Network Aging-associated Genes “shaken, not stirred” in a 3-Week Microgravity Simulation Human Study. bioRxiv. 2025 Jan 19. doi:10.1101/2025.01.19.633763

7. Kuznetsov NV, Vlasova DD, Kotikova AA, Tomilovskaya E, Ljubisavljevic M. Microgravity Impacts the Expression of Aging-Associated Candidate Gene Targets in the p53 Regulatory Network. Int J Mol Sci. 2025;26(22):11140. doi:10.3390/ijms262211140

8. European Physical Journal Plus. Radiation risk mitigation in human space exploration: a primer, a vision, and the state of the art. Eur Phys J Plus. 2026;141:100. doi:10.1140/epjp/s13360-025-07199-8

9. Kuznetsov NV, Almuzzaini B, Kritikou JS, Baptista MAP, Oliveira MMS, Keszei M, Snapper SB, Percipalle P, Westerberg LS. Nuclear Wiskott Aldrich syndrome protein co regulates T cell factor 1 mediated transcription in T cells. Genome Med. 2017;9:91.

10. Kuznetsov NV. Cell responses to simulated microgravity and hydrodynamic stress can be distinguished by using comparative transcriptomic analysis. bioRxiv. 2021. doi:10.1101/2021.12.31.474650

11. Kuznetsov NV. Cell Responses to Simulated Microgravity and Hydrodynamic Stress Can Be Distinguished by Comparative Transcriptomics. Int J Transl Med. 2022;2:364–86.

12. Hicks J, Olson M, Mitchell C, Juran CM, Paul AM. The Impact of Microgravity on Immunological States. ImmunoHorizons. 2023;7(10):670–82.

13. Lv H, Yang H, Jiang C, Shi J, Chen R, Huang Q, Shao D. Microgravity and immune cells. J R Soc Interface. 2023;20:20220869. doi:10.1098/rsif.2022.0869

14. Fava M, De Dominicis N, Forte G, Bari M, Leuti A, Maccarrone M. Cellular and Molecular Effects of Microgravity on the Immune System: A Focus on Bioactive Lipids. Biomolecules. 2024;14(4):446.

15. Buchheim JI, Feuerecker MF, Balsamo M, Vukich M, Van Walleghem M, Tabury K, et al. Monitoring functional immune responses with a cytokine release assay: ISS flight hardware design and experimental protocol for whole blood cultures executed under microgravity conditions. Front Physiol. 2024;14:1322852.

16. van den Nieuwenhof DWA, Moroni L, Chou J, Hinkelbein J. Cellular response in three dimensional spheroids and tissues exposed to real and simulated microgravity: a narrative review. NPJ Microgravity. 2024;10:95. doi:10.1038/s41526-024-00442-z

17. Wu F, Du H, Overbey E, Kim J, Makhijani P, Martin N, et al. Single-cell analysis identifies conserved features of immune dysfunction in simulated microgravity and spaceflight. Nat Commun. 2024;15(1):1995. doi:10.1038/s41467-023-42013-y

18. Martinez EM, Pelegrine AA, Holliday LS. Receptors Implicated in Microgravity-Induced Bone Loss. Receptors. 2023;2(1):14–28.

19. Chang TT, Walther I, Li CF, Boonyaratanakornkit J, Galleri G, Meloni MA, et al. The Rel/NF-κB pathway and transcription of immediate early genes in T cell activation are inhibited by microgravity. J Leukoc Biol. 2012;92(6):1133–45.

20. Hwang JR, Byeon Y, Kim D, Park SG. Recent insights of T cell receptor mediated signaling pathways for T cell activation and development. Exp Mol Med. 2020;52:750–61.

21. Boonyaratanakornkit JB, Cogoli A, Li CF, Schopper T, Pippia P, Galleri G, et al. Key gravity-sensitive signaling pathways drive T cell activation. FASEB J. 2005;19(14):2020–2.

22. Johansen KH, Golec DP, Thomsen JH, Schwartzberg PL, Okkenhaug K. PI3K in T cell adhesion and trafficking. Front Immunol. 2021;12:708908.

23. Di Luoffo M, Ben Meriem Z, Lefebvre P, Delarue M, Guillermet Guibert J. PI3K functions as a hub in mechanotransduction. Trends Biochem Sci. 2021;46(11):878–88.

24. Tauber S, Hauschild S, Paulsen K, Gutewort A, Raig C, Hürlimann E, et al. Signal transduction in primary human T lymphocytes in altered gravity during parabolic flight and clinostat experiments. Cell Physiol Biochem. 2015;35(3):1034–51.

25. Peterson AF, Ingram K, Huang EJ, et al. Systematic analysis of the MAPK signaling network reveals MAP3K driven control of cell fate. Cell Syst. 2022;13(11):885–94.e4.

26. Granet C, Boutahar N, Vico L, Alexandre C, Lafage-Proust MH. MAPK and SRC-Kinases Control EGR-1 and NF-κB Inductions by Changes in Mechanical Environment in Osteoblasts. Biochem Biophys Res Commun. 2002;293(5):1414–20.

27. Song D, Lian Y, Zhang L. The potential of activator protein 1 (AP-1) in cancer targeted therapy. Front Immunol. 2023;14:1224892.

28. Tan X, Xu A, Zhao T, Zhao Q, Zhang J, Fan C, et al. Simulated microgravity inhibits cell proliferation and activates MAPK pathway in human bone marrow mesenchymal stem cells. Life Sci Space Res. 2021;28:52–60.

29. Frigo DE, Tang Y, Beckman BS, et al. Mechanism of AP-1 mediated gene expression by select organochlorines through the p38 MAPK pathway. Carcinogenesis. 2004;25(2):249–61.

30. Zhang Y, Lu T, Wong M, Wang X, Stodieck L, Karouia F, et al. The MAPK pathway is differentially regulated in microgravity-exposed human T lymphocytes. FASEB J. 2020;34(S1):1–1.

31. Kuznetsov NV, Statsenko Y, Ljubisavljevic M. An update on neuroaging on Earth and in spaceflight. Int. J. Mol. Sci. 2025;26(4):1738.

32. Jeon H, Kim S, Park J, et al. Effects of simulated microgravity on colorectal cancer organoids growth and drug response. Sci Rep. 2024;14:25526. doi:10.1038/s41598-024-76737-8

33. Venkatraman S, Balasubramanian B, Thuwajit C, Meller J, Tohtong R, Chutipongtanate S. Targeting MYC at the intersection between cancer metabolism and oncoimmunology. Front Immunol. 2024;15:1324045.

34. Kim H, Shin Y, Kim DH. Mechanobiological Implications of Cancer Progression in Space. Front Cell Dev Biol. 2021;9:740009.

35. Topal U, Zamur C. Microgravity, Stem Cells, and Cancer: A New Hope for Cancer Treatment. Stem Cells Int. 2021;2021:5566872.

36. Becker JL, Souza GR. Using space-based investigations to inform cancer research on Earth. Nat Rev Cancer. 2013;13(5):315–27.

37. Vora PM, Prabhu S. Exploring the influence of microgravity on chemotherapeutic drug response in cancer: Unveiling new perspectives. J Cell Mol Med. 2024;28(9):e18342.

38. Grimm D, Corydon TJ, Sahana J, González Torres LF, Kraus A, Marchal S, et al. Recent studies of the effects of microgravity on cancer cells and the development of 3D multicellular cancer spheroids. Stem Cells Transl Med. 2025;14(3):szaf008.

39. Graf J, Schulz H, Wehland M, Corydon TJ, Sahana J, Abdelfattah F, et al. Omics studies of tumor cells under microgravity conditions. Int J Mol Sci. 2024;25(2):926.

40. Corydon TJ, Schulz H, Richter P, Strauch SM, Böhmer M, Ricciardi DA, et al. Current Knowledge about the Impact of Microgravity on Gene Regulation. Cells. 2023;12(7):1043.

41. Blutt SE, Crawford SE, Ramani S, Zou WY, Estes MK. The impact of microgravity on intestinal stem cells and epithelial regeneration. Stem Cells Dev. 2022;31(15–16):447–58.

42. Schuhwerk H, Kleemann J, Gupta P, Stemmler MP, Brabletz T, Brabletz S. The EMT transcription factor ZEB1 governs a fitness promoting but vulnerable DNA replication stress response. Cell Rep. 2022;41(11):111819.

43. Chen Z, Luo Q, Lin C, Kuang D, Song G. Simulated microgravity inhibits osteogenic differentiation of mesenchymal stem cells through downregulating ZEB1. FASEB J. 2019;33(1):891–904.

44. Ulbrich C, Westphal K, Pietsch J, Winkler HD, Leder A, Bauer J, et al. Characterization of human chondrocytes exposed to simulated microgravity. Cell Physiol Biochem. 2010;26(3):379–90.

45. Quang MT, Nguyen MN. The potential of microRNAs in cancer diagnostic and therapeutic strategies: a narrative review. J Basic Appl Zool. 2024;85:7.

46. Girardi C, De Pittà C, Casara S, Calura E, Romualdi C, Celotti L, et al. Integrated analysis of microRNA and mRNA expression profiles in peripheral blood lymphocytes of astronauts. Sci Rep. 2020;10:8175.

47. Vidyasekar P, Shyamsunder P, Arun R, Santhakumar R, Kapadia NK, Kumar R, et al. Genome wide expression profiling of cancer cell lines cultured in microgravity reveals significant dysregulation of cell cycle and microRNA gene networks. PLoS One. 2015;10(8):e0135958.

48. Mangiapane G, Nicotera G, Peresson M, Pignatelli A, Bizzarri M, Cucina A, et al. MicroRNA expression profiling in simulated microgravity-treated human colon cancer cells. Int J Mol Sci. 2022;23(15):8341.

49. Zhao T, Li R, Tan X, Zhang J, Fan C, Zhao Q, et al. MicroRNA-mediated regulation of cell cycle and apoptosis in microgravity-exposed cancer cells. Front Oncol. 2021;11:678912.

50. Prasad B, Grimm D, Strauch SM, Erzinger GS, Corydon TJ, Lebert M, et al. Influence of Microgravity on Apoptosis in Cells, Tissues, and Other Systems In Vivo and In Vitro. Int J Mol Sci. 2020;21(24):9373.

51. Fukazawa T, Tanimoto K, Shrestha L, Imura T, Takahashi S, Sueda T, et al. Simulated microgravity enhances CDDP induced apoptosis signal via p53 independent mechanisms in cancer cells. PLoS One. 2019;14(7):e0219363.

52. Hao Q, Chen J, Lu H, Zhou X. The ARTS of p53 dependent mitochondrial apoptosis. J Mol Cell Biol. 2022;14(10):mjac074.

53. Zhou Z, Arroum T, Luo X, Kang R, Lee YJ, Hüttemann M, Song X. Diverse functions of cytochrome c in cell death and disease. Cell Death Dis. 2024;15(8):603.

54. Wang Y, Li X, Zhang Y, et al. PIEZO1 Mediates Apoptosis of Endothelial Cells via Enhancing HMGA2 Expression Under Simulated Microgravity. Int J Mol Sci. 2026;27(3):1425. doi:10.3390/ijms27031425

55. Kleinsimon S, Longmuss E, Rolf J, Jäger S, Eggert A, Delebinski C, Seifert G. GADD45A and CDKN1A are involved in apoptosis and cell cycle modulatory effects: implications for stress induced growth arrest. Sci Rep. 2018;8:10072.

56. Kanda K, Yamauchi K, Kurabayashi T, Inoue R. Simulated microgravity enhances apoptosis in vascular endothelial cells via the mitochondrial pathway. J Gravit Physiol. 2020;27(1):25–32.

57. Li N, Wang C, Sun S, Zhang C, Lü D, Chen Q, et al. Microgravity-induced apoptosis and mitochondrial dysfunction in human lymphoblastoid cells. Apoptosis. 2019;24(7–8):619–32.

58. Blaber EA, Dvorochkin N, Lee C, Alwood JS, Yousuf R, Pianetta P, et al. Microgravity induces pelvic bone loss through osteoclastic activity, osteocytic osteolysis, and osteoblastic cell cycle inhibition by CDKN1A/p21. PLoS One. 2015;10(8):e0133937.

59. Humayun A, Fornace AJ Jr. GADD45 in stress signaling, cell cycle control, and apoptosis. Adv Exp Med Biol. 2022;1360:1–22.

60. Blaber EA. Osteogenic Differentiation of Somatic Stem Cells in Space: A Study Investigating the Role of CDKN1a/p21 on Mesenchymal Stem Cell Proliferation, Differentiation, and Regeneration in Microgravity. NASA OSDR. Experiment OS-753.

61. Dai ZQ, Wang R, Ling SK, Wan YM, Li YH. Simulated microgravity inhibits the proliferation and osteogenesis of human bone marrow mesenchymal stem cells by affecting the cell cycle. Biochem Biophys Res Commun. 2020;527(3):703–9.

62. Poon CE, Madueke IC, Stodieck LS, Louis SA, Blaber EA. Identification of CDKN1a/p21 as a key regulator of mesenchymal stem cell proliferation in microgravity. Stem Cells Dev. 2021;30(12):619–31.

63. Lane, D.P. p53, guardian of the genome. Nature. 1992, 358, 6381.

64. Zaĭchuk, T.A.; Kuznetsov, N.V.; Osovskaia, V.S.; Kopnin, B.P.; Chumakov, P.M. Specific DNA-binding properties of oncoprotein p53 in human tumor cells. Dokl. Akad. Nauk. 1993; 330:379–382.

65. Zaichuk, T.A.; Kuznetsov, N.V.; Ossovskaya, V.S.; Kopnin, B.J.; Chumakov, P.M. Specific DNA-binding properties of p53 oncoprotein from human tumor cells. Proc. Russ. Acad. Sci. 1993; 330:386–389.

66. Kuznetsov, N.V. Analysis of DNA-Binding Properties of Protein P53. 1995. Available online: www.dslib.net/molekula/analiz-dnk-svjazyvajuwih-svojstv-belka-r53.html.

67. Kuznetsov NV, Andersson P, Gradin K, Stein PV, Dieckmann A, Pettersson S, Hanberg A, Poellinger L. The dioxin/aryl hydrocarbon receptor mediates downregulation of osteopontin gene expression in a mouse model of gastric tumourigenesis. Oncogene. 2005 Apr;24(19):3216–22.

68. Kondratov, R.V.; Kuznetsov, N.V.; Pugacheva, E.N.; Almazov, V.P.; Prasolov, V.S.; Kopnin, B.P.; Chumakov, P.M. Functional heterogeneity of p53-responsive elements. Mol. Biol. 1996; 30:613–620.

69. Kondratov, R.V.; Pugacheva, E.N.; Kuznetsov, N.V.; Prasolov, V.S.; Kopnin, B.P.; Chumakov, P.M. The human adenosine deaminase gene contains a p53-responsive element. Dokl. Akad. Nauk. 1996; 346: 260–262.

70. Chumakov, P.M.; Pugacheva, E.N.; Ossovskaya, V.S.; Kondratov, R.V.; Kuznetsov, N.V.; Ivanov, A.V.; Prassolov, V.S.; Mazo, I.A.; Gudkov, A.V.; Kopnin, B.P. Functions of the p53 involved in control of malignant transformation. In Cancer and the Cell Cycle; ISREC Press: Lausanne, Switzerland, 1996.

71. Kondratov, R.V.; Kuznetsov, N.V.; Pugacheva, E.N.; Almazov, V.P.; Prasolov, V.S.; Chumakov, P.M.; Kopnin, B.P. Functional heterogeneity of p53-responsive elements. Mol. Biol. 1996; 30 (Suppl. 2):363–367.

72. Chumakov, P.M. Versatile functions of p53 protein in multicellular organisms. Biochemistry 2007; 72:1399–1421.

73. Link W, Ferreira BI. FOXO Transcription Factors: A Brief Overview. Methods Mol Biol. 2024; 2871:1–8.

74. Kaiser FMP, Janowska I, Menafra R, de Gier M, Korzhenevich J, Pico Knijnenburg I, et al. IL-7 receptor signaling drives human B cell progenitor differentiation and expansion. Blood. 2023;142(13):1113–30.

75. Waickman AT, Keller HR, Kim TH, et al. The Cytokine Receptor IL-7Rα Impairs IL-2 Receptor Signaling and Constrains the In Vitro Differentiation of Foxp3+ Treg Cells. iScience. 2020;23(8):101421.

76. Brunet A, Datta SR, Greenberg ME. Transcription-dependent and -independent control of neuronal survival by the PI3K-Akt-FOXO signaling pathway. Curr Opin Neurobiol. 2021;11(3):297–305.

77. Asadi YY, Moundounga RK, Chakroborty AC, et al. FOXOs and their roles in acute and chronic neurological disorders. Front Mol Biosci. 2025;12:1538472.

78. Smith JK. Osteoclasts and Microgravity. Life (Basel). 2020;10(9):207.

79. Smith CI, Burger JA. Bruton tyrosine kinase in B cell development and primary immunodeficiency: mechanisms and clinical implications. Nat Rev Immunol. 2021;21(6):345–60.

80. Cameron B, Zaheer SA, Dominguez Villar M. Control of CD4+ T cell differentiation and function by PI3K isoforms. In: PI3K and AKT Isoforms in Immunity. Curr Top Microbiol Immunol. 2022;436:197–216.

81. Phelan MA, Gianferante DM, Cage CL, Urbanski ML, Basner M, Smith SM, et al. The Impact of Spaceflight on Humoral Immunity and Vaccine Response. J Immunol. 2022;208(1):12–21.

82. Crucian B, Stowe R, Mehta S, Quiriarte H, Pierson D, Sams C. Alterations in adaptive immunity persist during long-duration spaceflight. NPJ Microgravity. 2015;1:15013.

83. Krüger M, Melnik D, Kopp S, Buken C, Sahana J, Bauer J, et al. Monitoring of T-cell activation and differentiation in microgravity. Int J Mol Sci. 2019;20(23):5925.

84. Takegahara N, Kim H, Choi Y. Unraveling the intricacies of osteoclast differentiation and maturation: insight into novel therapeutic strategies for bone destructive diseases. Exp Mol Med. 2024;56(4):789–98.

85. Man J, Graham T, Squires Donelly G, Laslett AL. The effects of microgravity on bone structure and function. NPJ Microgravity. 2021;7:12.

86. Zhang X, Xue T, Hu Z, Gao XG, Li L, Wang Y, et al. Bioinformatic analysis of the RNA expression patterns in microgravity induced bone loss. Front Genet. 2022;13:985025.

87. Tamma R, Colaianni G, Camerino C, Di Benedetto A, Greco G, Strippoli M, et al. Microgravity during spaceflight directly affects osteoclastogenesis by increasing RANKL/OPG ratio. FASEB J. 2022;36(5):e22312.

88. Nabavi N, Khandani A, Camirand A, Harrison RE. Effects of microgravity on osteoclast differentiation and activity. J Cell Biochem. 2021;122(8):908–20.

89. Saxena R, Pan G, McDonald JM. Osteoclast and osteoblast responses to modeled microgravity: a systems biology analysis. NPJ Microgravity. 2020;6:17.

90. Calderwood DA, Shattil SJ, Ginsberg MH. Integrins and Actin Filaments: Reciprocal Regulation of Cell Adhesion and Signaling. J Biol Chem. 2000;275(30):22607–10.

91. Tauber S, Lauber BA, Paulsen K, Layer LE, Lehmann M, Hauschild S, et al. Cytoskeletal stability and metabolic alterations in primary human macrophages in long term microgravity. PLoS One. 2017;12(4):e0175599.

92. López Garzón NA, Pinzón Fernández MV, Saavedra JT, Nati Castillo HA, Arias Intriago M, Salazar Santoliva C, Izquierdo Condoy JS. Microgravity and Cellular Biology: Insights into Cellular Responses and Implications for Human Health. Int J Mol Sci. 2025;26(7):3058.

93. Gardiner J. Cytoskeletal Tensegrity in Microgravity. Life (Basel). 2021;11(10):1091.

94. Vorselen D, Roos WH, MacKintosh FC, Wuite GJ, van Loon JJ. The role of the cytoskeleton in sensing changes in gravity by mechanosensitive channels. FASEB J. 2020;34(2):2342–57.

95. Gerasimčik N, He M, Dahlberg CI, Kuznetsov NV, Severinson E, Westerberg LS. The small Rho GTPases Rac1 and Rac2 are important for T cell independent antigen responses and for suppressing switching to IgG2b in mice. Front Immunol. 2017;8:1264.

96. Louis F, Deroanne C, Nusgens B, Vico L, Guignandon A. RhoGTPases as key players in mechanotransduction during microgravity exposure. Cell Mol Life Sci. 2022;79(3):152.

97. Thiel CS, Hauschild S, Huge A, Tauber S, Lauber BA, Polzer J, et al. Dynamic gene expression response to altered gravity in human T cells. Sci Rep. 2021;11:16534.

98. Kuznetsov NV. Search for genes of T cell response to simulated weightlessness in vitro using comparative transcriptomic analysis. In: XLVII Academic Readings on Cosmonautics. 2023. p. 42–4.

99. Paulsen K, Tauber S, Gölz N, Layer LE, Simmet T, Fink B, et al. The cytoskeleton and its role in T cell migration and activation under microgravity conditions. Cell Commun Signal. 2020;18(1):112.

100. Stratis D, Morin M, Gallois L, et al. The Characteristic Response of the Human Leukocyte Transcriptome to 60 Days of Bed Rest and to Reambulation. Med Sci Sports Exerc. 2023. doi:10.1249/MSS.0000000000003085

101. Schmidt MA, Goodwin TJ. Personalized medicine in human space flight: using Omics based analyses to develop individualized countermeasures. Metabolomics. 2013;9(6):1134–1156. doi:10.1007/s11306-013-0556-3

102. Guo Y, Waisberg E, Ong J, Kumar R, Lee AG. A future of personalized medicine for astronauts: Considering genetic variability and biologic sex-based differences in space medicine. Acta Astronautica. 2025;228:527–531. doi:10.1016/j.actaastro.2024.12.035

